# A G-lectin Receptor Kinase is a Negative Regulator of Arabidopsis Immunity Against Root-Knot Nematode *Meloidogyne incognita*

**DOI:** 10.1101/2021.09.07.459316

**Authors:** Dongmei Zhou, Damaris Godinez-Vidal, Jiangman He, Marcella Teixeira, Jingzhe Guo, Lihui Wei, Jaimie M. Van Norman, Isgouhi Kaloshian

**Author notes:** **Author for correspondence:** Isgouhi Kaloshian, Department of Nematology, University of California Riverside, CA, USA. These authors contributed equally (Dongmei Zhou, Damaris Godinez-Vidal). Senior author (Isgouhi Kaloshian). **Author contributions:** I.K., J.M.V.N. and L.W. planned and designed the experiments; D.Z., J.H., D.G.-V., M.T., J.G. and J.M.V.N. performed the experiments; I.K. and J.M.V.N. supervised the experiments and analyzed the data; I.K. wrote the manuscript with contributions from all authors. I.K. agrees to serve as the author responsible for contact and ensures communication. Jiangman He: Department of Botany and Plant Sciences, University of California Riverside, CA 92521, USA. Conflicts of interest statement. The authors declare no conflicts of interest.

## Abstract

Root-knot nematodes (*Meloidogyne* spp., RKN) are responsible for extensive crop losses worldwide. For infection, they penetrate plant roots, migrate between plant cells, and establish feeding sites, known as giant cells, in the root pericycle. Previously, we found that nematode perception and early plant responses were similar to those for microbial pathogens and require the BAK1 co-receptor in *Arabidopsis thaliana* and tomato. To identify additional receptors involved in this process, we implemented a reverse genetic screen for resistance or sensitivity to RKN using Arabidopsis T-DNA alleles of genes encoding transmembrane receptor-like kinases. This screen identified a pair of allelic mutations with enhanced resistance to RKN in a gene we named *ENHANCED RESISTANCE TO NEMATODES 1 (ERN1). ERN1* encodes a G-type lectin receptor kinase (G-LecRK) with a single pass transmembrane domain. Further characterization showed that *ern1* mutants displayed stronger activation of MAP kinases, elevated levels of the defense marker *MYB51*, and enhanced H_2_0_2_ accumulation in roots upon RKN elicitor treatments. Elevated *MYB51* expression and ROS burst were also observed in leaves of *ern1* mutants upon flg22 treatment. Complementation of *ern1.1* with 35S- or native promotor-driven *ERN1* rescued the RKN infection and enhanced defense phenotypes. Taken together, our results indicate that *ERN1* is an important negative regulator of immunity.

**One sentence summary:** A plasma membrane localized G-lectin receptor kinase acts as a negative immune regulator by interfering with defense responses activated by nematode and microbial elicitors.

## Introduction

The majority of plant parasitic nematodes (PPN) are soil dwelling organisms that use a specialized mouth part or stylet to penetrate root tissues and establish an intimate relationship with their hosts. Plant-nematode signaling starts even before nematode penetration, as nematodes are attracted by root diffusates dispersed in the soil (Goverse and Smant, 2014). After finding their host, different species of PPN deploy specific strategies to penetrate plant tissues and initiate feeding. Root-knot nematodes (*Meloidogyne* spp., RKN) are the most devastating group of PPN causing great crop losses worldwide (Jones et al., 2013; Kaloshian and Teixeira, 2019). RKN are sedentary endoparasites penetrate plant roots behind the root tip and migrate between cells until they reach the vascular cylinder, where they establish a specialized feeding site (Wyss et al., 1992; Goverse and Smant, 2014). RKN secretions, mainly originating from esophageal glands, contribute to massive reprograming of plant cell processes, including cell division. The outcome is the development of enlarged and multinucleated cells termed giant cells, resulting from karyokinesis without cytokinesis. Each nematode forms about six to eight giant cells surrounded by enlarged neighboring endodermis and cortical cells, forming a root gall. This feeding site acts as a nutrient sink, nourishing the RKN during its entire life cycle. Once the feeding site is established, RKNs become sedentary and as females mature, they become pearl-shaped with the reproductive organs occupying most of their body. Most RKN species reproduce parthenogenetically. A gelatinous sac develops at its posterior end, protruding onto the surface of the root, where a large number of eggs are laid.

Plant defense responses are triggered by elicitors derived from microbes. General elicitors, or microbe-associated-molecular patterns (MAMP), are perceived by cell surface localized pattern recognition receptors (PRRs) and trigger an immune response known as pattern-triggered immunity (PTI) (Yu et al., 2017). A number of PRRs have been identified and among them is the well characterized receptor FLS2 (FLAGELLIN SENSING2). *Arabidopsis* FLS2 recognizes a highly conserved stretch of 22 amino acids, flg22, present on the N-terminus of bacterial flagellin, which acts as a molecular glue that brings together FLS2 and the co-receptor BAK1/SERK3 (BRI1-ASSOCIATED KINASE1/SOMATIC EMBRYOGENESIS RECEPTOR KINASE3), to elicit downstream signaling events, including phosphorylation events, transcriptional reprograming, callose deposition, and ROS burst (Felix et al., 1999; Gomez-Gomez et al., 1999; Zipfel et al., 2004; Chinchilla et al., 2007). Notably, flg22-mediated defense elicitation is followed by transcriptional upregulation of *FLS2* and *BAK1*, as well as upregulation of negative regulators of immunity, such as *PBL13* (*AvrPphB SUSCEPTIBLE1-LIKE13*) (Lin et al., 2015). Additionally, after elicitation of defense responses, FLS2 is internalized from the plasma membrane to internal vesicles, likely for degradation (Salomon and Robatzek, 2006; Ben Khaled et al., 2015).

Recently, processes involved in root perception of nematodes and early responses during nematode penetration have been unveiled. It was shown that RKN and cyst nematodes (CNs) infective-stage juveniles are perceived by plant roots during their root migration phase, similar to the perception of microbial pathogens in above ground tissues (Mendy et al., 2017; Teixeira et al., 2016). RKN perception required the co-receptor BAK1/SERK3 as *SERK3*-silenced tomato plants displayed enhanced susceptibility to RKN (Peng and Kaloshian, 2014). BAK1/SERK3 is a coreceptor of multiple MAMPs coordinating perception with diverse PRRs to activate PTI. Enhanced susceptibility to both RKN and CN were also reported in the *Arabidopsis bak1*-*5* mutant (Mendy et al., 2017; Teixeira et al., 2016). Interestingly, a leucine-rich repeat (LRR) serine/threonine kinase, NILR1 (NEMATODE-INDUCED LRR-RLK 1), was shown to have similar characteristics as microbial PRRs and, therefore, could be a receptor of an, as yet, unidentified nematode-associated molecular pattern (NAMP) (Mendy et al., 2017). In addition, mutants in *Arabidopsis* BIK1 (BOTRYTIS-INDUCED KINASE1), which is associates with and is phosphorylated by BAK1, and the double mutant of the RBOH D/F (RESPIRATORY BURST NADPH OXIDASE HOMOLOG), which are phosphorylated by BIK1, also displayed enhanced susceptibility to RKN (Lu et al., 2010; Zhang et al., 2010; Lin et al 2014; Li et al., 2014; Kadota et al., 2014; Teixeira et al., 2016). Taken together, these findings indicate that canonical PTI signaling is involved in RKN perception.

Despite the great potential of the broad and lasting defense mediated by PTI responses, nematode-related immunity research has largely focused on characterization of resistance (R) gene-mediated defense and nematode effectors (Gheysen and Mitchum, 2011; Goverse and Smant, 2014; Kaloshian and Teixeira, 2019). Nevertheless, a few investigations have characterized transcriptional changes in response to nematode infection at early time points. One of the first descriptions of gene expression changes found eight differentially expressed genes in response to RKN both in susceptible and resistant tomato (*Solanum lycopersicum*) plants only 12h after inoculation, when nematodes were still migrating through plant tissues (Lambert et al., 1999). Interestingly, the transcriptome of tomato roots collected 24h after RKN inoculation showed consistently more genes upregulated than downregulated among those that were differentially expressed, including defense-related genes (Bhattarai et al., 2008). More recently, using reporter GUS lines, activation of defense-related genes was shown in *Arabidopsis* roots exposed to crude RKN extracts providing a novel tool for evaluation of defense responses in the absence of wound-induced responses due to RKN penetration (Teixeira et al., 2016).

To identify additional immune receptors involved in the PTI response against RKN, we used a reverse genetics approach to screen *Arabidopsis* T-DNA insertion alleles of receptor-like kinases (RLKs). Instead of a receptor acting as a positive regulator of immunity, we identified a novel negative regulator of RKN immunity. Here we present the characterization of a negative regulator of RKN immunity encoded by a lectin receptor kinase.

## RESULTS

### Mutation of the Arabidopsis RLK ERN1 results in resistance to RKN

To identify putative receptors involved in immunity against RKN, we screened homozygous *Arabidopsis* T-DNA insertional mutants in genes encoding transmembrane receptor-like kinases for resistance or sensitivity to RKN. At least two mutant alleles for each gene were screened (Supplemental Table S1). Among the mutants, those with T-DNA insertions in At1g61550, SALK_128729 and SAIL_63_G02, displayed significantly fewer galls on roots indicating enhanced resistance to RKN infection (Figure 1A). Accordingly, the locus was named *ENHANCED RESISTANCE TO NEMATODES* (*ERN1*) and the mutants designated *ern1.1* (SALK_128729) and *ern1.2* (SAIL_63_G02) (Figure 1A).

**Figure 1.**
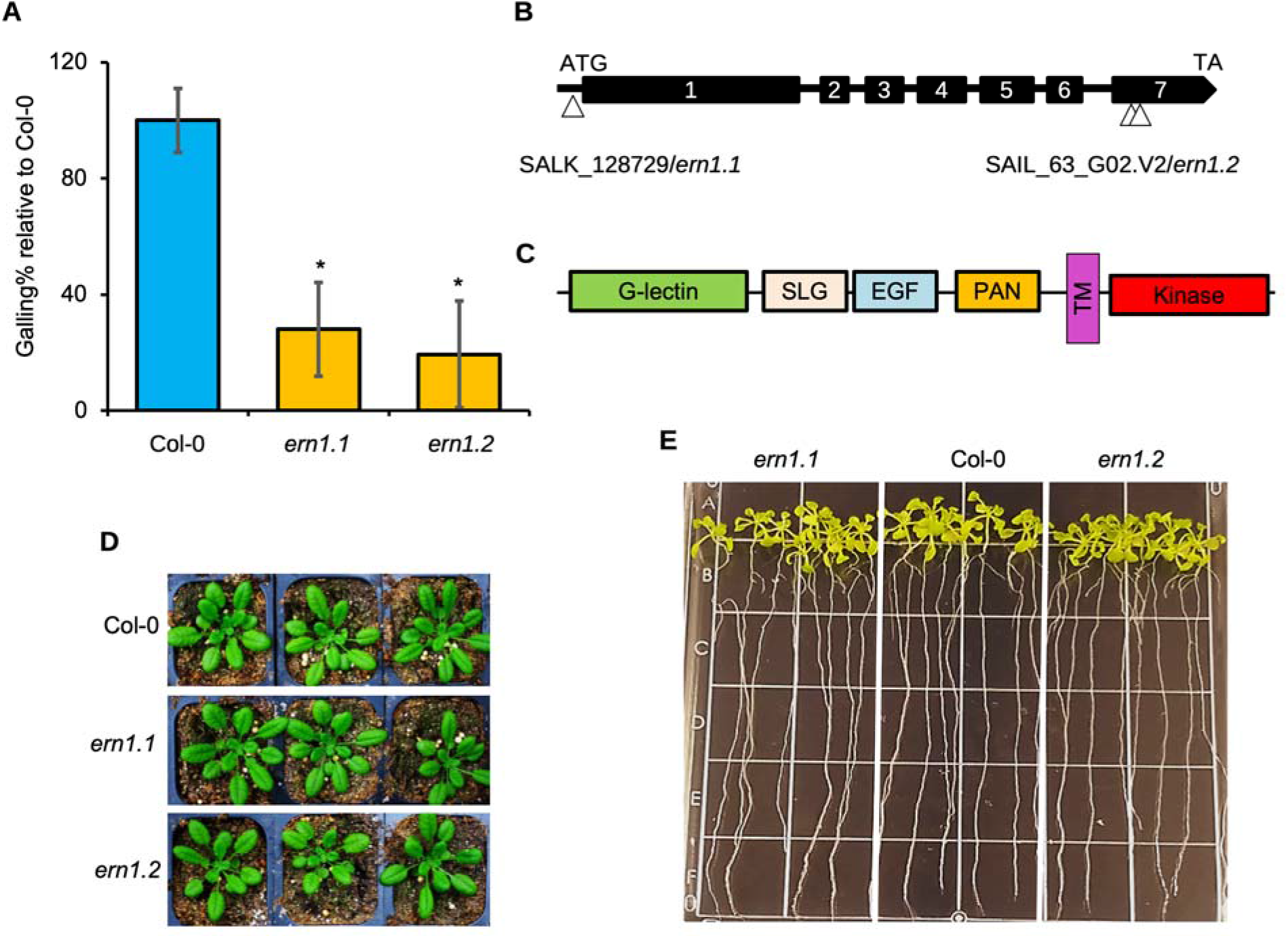
Arabidopsis *ern1* mutant resistance phenotype, *ERN1* gene/protein structure, and *ern1* morphological phenotypes. (A) Nematode infection phenotype of Arabidopsis wildtype Col-0, *ern1.1* (SALK_128729) and *ern1.2* (SAIL_63_G02) T-DNA mutants, grown on Gamborg media and infected with 100 J2. Bars are percentage of galls on roots normalized to wildtype ±SE (n=75 per genotype). Asterisks indicate significance difference (*P* < 0.001, ANOVA). (B) SALK_128729/*ern1.1* has an insertion in the first exon while the SAIL_63_G02/*ern1.2* mutant has at least two insertions in the seventh exon of the gene. (C) ERN1 predicated protein structure: S-locus glycoprotein (SLG), epidermal growth factor-like (EGF), and a plasminogen-apple-nematode (PAN), transmembrane (TM) domains. (D) Shoot growth phenotype of 3-week-old Arabidopsis plants grown under 16h photoperiod at 22°C. (E) Root growth phenotype of 3-week-old Arabidopsis plants grown on MS agar plates under 12h photoperiod at 22°C.

### Characterization of *ERN1* and its mutant alleles

Sequence analysis indicated that the *ERN1* genomic sequence is 3,629 base pairs (bp) with 7 exons and a cDNA of 2,982 bp (Figure 1B). Protein domain examination using Interpro revealed that *ERN1* encodes a G-lectin receptor kinase (G-LecRK) with a G-type lectin domain, a single transmembrane domain, and an intracellular serine/threonine kinase domain (Figure 1C). In addition, ERN1 contains a S-locus glycoprotein, an epidermal growth factor-like (EGF), and a plasminogen-apple-nematode (PAN) extracellular domains (Figure 1C). Phylogenetic analysis of Arabidopsis G-LecRKs classified *ERN1* as G-LecRK-VIII.8 (Teixeira et al., 2018).

The T-DNA insertions in the *ern1.1* and *ern1.2* alleles were predicted to be in the first and seventh exons, respectively. We confirmed the locations of the T-DNA insertions in both mutants by PCR and sequencing. This analysis revealed that the insertion in *ern1.1* is located 49 bp upstream of *ERN1* start codon, while *ern1.2* has a deletion of 61bp and introduction of a premature stop codon in the predicted kinase domain (Figure 1B). In addition, our results indicated that *ern1.2* has at least two T-DNA insertions in tandem and reverse orientation (Figure 1B and Supplemental Figure S1A).

Assessing the expression of *ERN1* in these mutants indicated that *ern1.1* is a null allele as no *ERN1* transcripts were detected in this mutant, while a reduction in full length transcript and a deletion are detected in *ERN1* transcripts in *ern1.2* (Supplemental Figure S1B). The enhanced nematode resistance in the *ern1* mutants lead us to investigate whether the resistance phenotype is due to morphological changes in plant or root growth phenotypes. Wild type (Col-0), *ern1.1*, and *ern1.2* mutants were grown in soil under short (12h/12h L/D) and long (16h/8h L/D) day photoperiods at 22°C and under short day-length at 19°C. No difference in above ground growth morphology was detected between Col-0 and either of the *ern1* mutants under all tested conditions (Figure 1D; Supplemental Figure S2). Measurements of rosette leaf dimensions also did not reveal any leaf growth changes in the *ern1* mutants compared to WT (Supplemental Figure S3). Since RKNs infect behind root tips and alterations in root development may affect nematode infection rates, we investigated root growth in the *ern1* mutants. Seedlings grown on agar plates were evaluated for root growth over a 4-day period (days 4-8) and lateral root capacity at 8 days. No difference was detected in root growth rate or lateral root capacity between WT and the *ern1* alleles (Figure 1E and Supplemental Figure S4).

### The *ern1* mutants display higher levels of H_2_0_2_ accumulation and upregulation of immune genes induced by RKN elicitors

The enhanced resistance displayed by the *ern1* mutants and lack of obvious root growth phenotypes suggests enhanced immunity in these mutants. To evaluate enhanced immune responses in roots, we treated the roots of *ern1* and wild type plants with RKN NemaWater, which is nematode-free water generated by incubating newly emerged infective-stage juveniles in water overnight. NemaWater from *M. incognita* and the cyst nematode *Heterodera schachtii* have been shown to trigger immune responses in different plant species, including *Arabidopsis* (Mendy et al 2017). NemaWater or water-treated roots were stained with immunohistochemical 3,3’diaminobenzidine (DAB) to evaluate H_2_0_2_ accumulation detected as a brown-stained precipitate. While control roots treated with water did not display any color change, both *ern1.1* and *ern1.2* mutant roots treated with NemaWater consistently showed more intense dark brown staining compared to WT roots (Figure 2A). Previously, we found that crude extracts from infective juveniles also triggered immune responses in roots, including activation of the transcription factor *MYB51* (Teixeira et al., 2016). We wondered whether nematode eggs, which contain different embryonic stages as well as first and second stage juveniles, could also trigger similar responses as NemaWater. We first evaluated the ability of nematode eggs to trigger *MYB51* expression in *Arabidopsis* roots. *Arabidopsis* plants expressing the *MYB51pro::GUS* reporter were treated with nematode egg extract and evaluated for GUS activity. Infective-stage juvenile extract was used as a positive control. GUS activity was detected in roots of *MYB51pro::GUS* reporter lines treated with either egg extract or infective juvenile extract indicating nematode egg extract is able to trigger immune responses similar to that from infective juveniles (Supplemental Figure S5). Similarly, *ern1* mutant roots treated with nematode egg extracts and stained with DAB also showed more intense brown staining in the roots compared to WT confirming increased H_2_0_2_ accumulation (Figure 2A).

**Figure 2.**
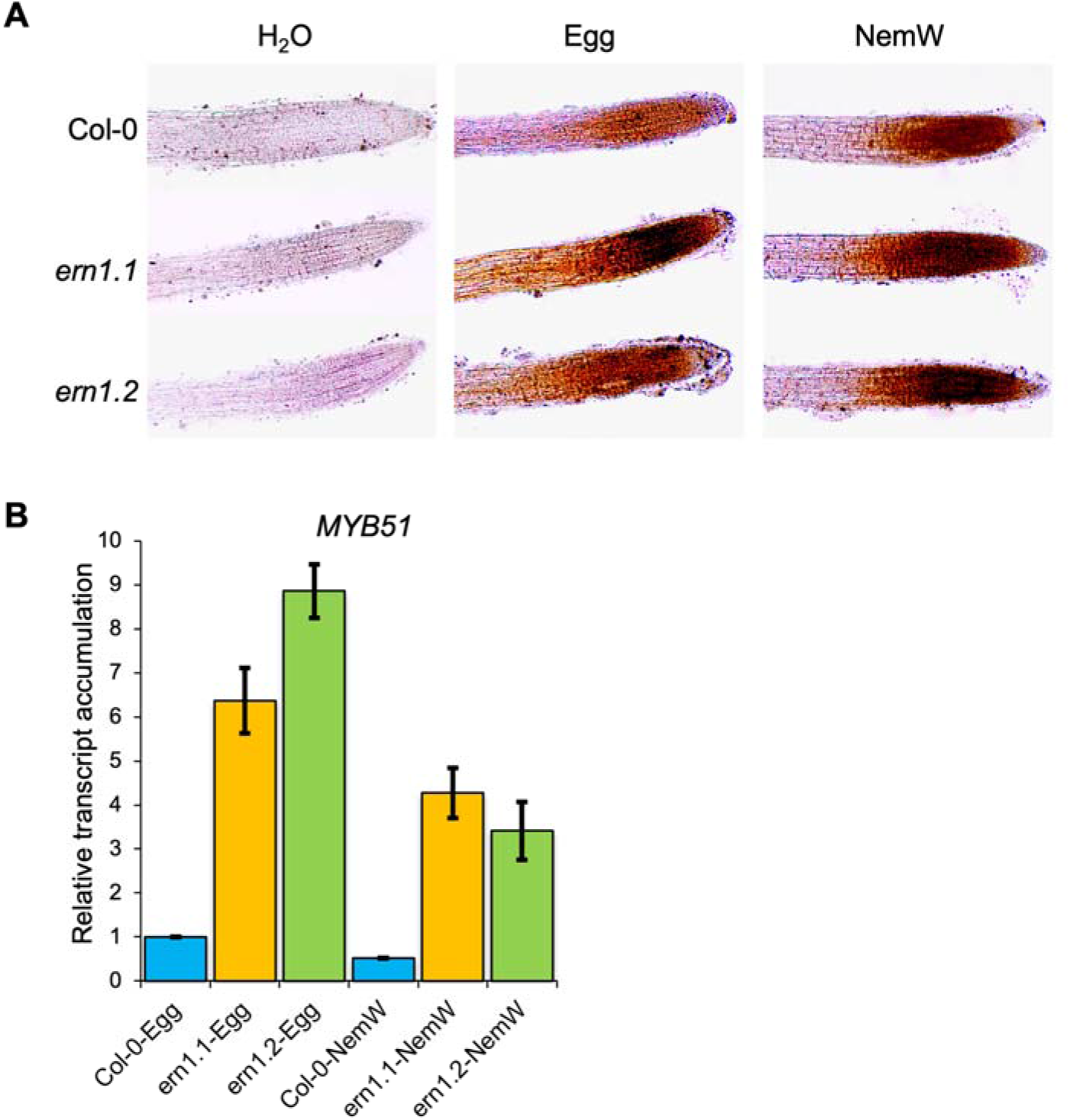
Root-knot nematode (RKN) egg extract and excretions induce enhanced immune responses in *ern1* roots. (A) DAB stained roots of 5-day-old Arabidopsis Col-0, *ern1.1*, and *ern1.2* seedlings treated with RKN egg extract (Egg) or Nemawater (NemW) for 1 h. The experiment was performed four times with similar results. (B) *MYB51* expression in roots of 5-day-old Arabidopsis Col-0, *ern1.1*, and *ern1.2* mutants treated with egg extract or NemW for 1 h and used for RNA extraction. Gene expression analysis was performed by qPCR and normalized to *UBQ22*. Bars are means of three biological replicates ± SE.

To quantitatively evaluate the enhanced resistance of *ern1* mutants, roots of *ern1* mutants and WT were treated with nematode egg extracts, NemaWater, or water and were used to examine *MYB51* expression by qPCR. Both nematode egg extracts and NemaWater induced *MYB51* transcripts levels to higher levels in the *ern1* mutant roots compared to WT (Figure 2B). Together these results suggest that enhanced resistance of *ern1* mutants may be attributed increased H_2_0_2_ accumulation and strong induction of *MYB51* expression, which are likely induced by different RKN elicitors.

### Egg-derived elicitor(s) activate MAP kinases at higher levels in *ern1* mutants

The mitogen-activated protein kinases MPK3 and MPK6 are activated upon perception of many MAPMs and elicitors (Asai et al., 2002). Given the enhanced resistant phenotype of *ern1* mutants, we explored whether nematode egg-derived elicitors induce MPK3/6 activity and if differential levels of kinase activity could be detected in *ern1* mutants. Since MPK3/6 activity is known to be an early and transient response, we performed a time-course experiment collecting tissues early after the egg extract treatment of *ern1* and WT roots and performed immunoblot analysis using anti-phospho-p42/44 antibody. Egg extract treatment resulted in rapid activation of MPK3/6, and more importantly, stronger activation was detected in *ern1* mutants compared to WT (Figure 3). As expected, MPK3/6 activation was transient reaching to non-detectable levels by 2 hours after egg extract treatment. These results further confirm that the egg-derived elicitors behave like the microbial elicitors.

**Figure 3.**
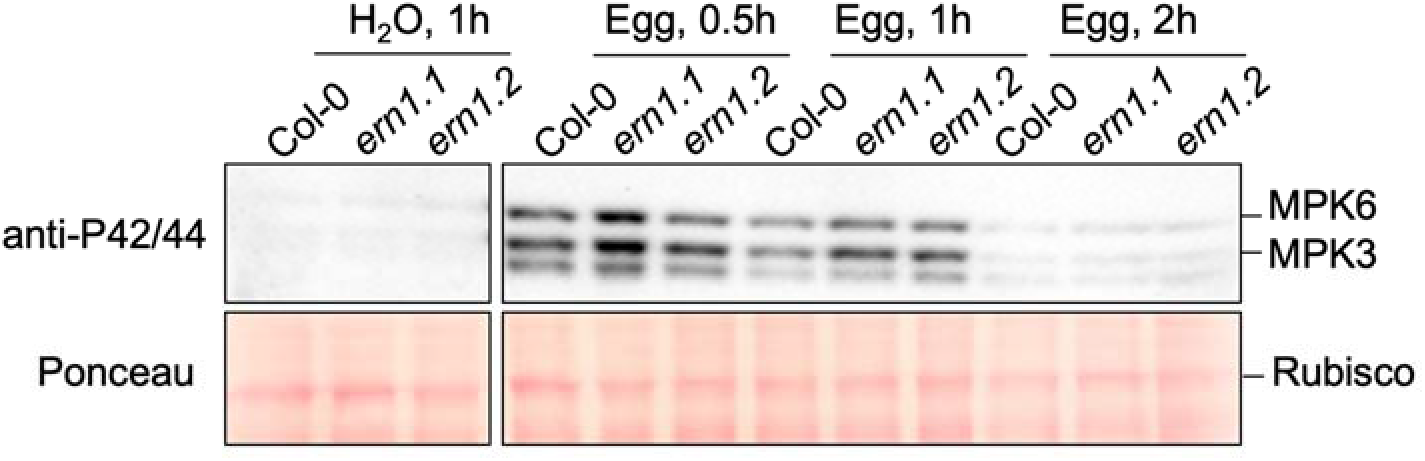
Root-knot nematode (RKN) egg extract-induced MAPK activation is enhanced in *ern1* roots. Roots of 12-day-old Arabidopsis Col-0, *ern1.1*, and *ern1.2* seedlings were treated with RKN egg extract (Egg) for 0.5h, 1 h, and 2h or water control for 1h and used for immunoblot analysis. Activated MAPKs were detected using anti-phospho-p42/44 MAPK antibody. Ponceau stained Rubisco was used for equal loading control.

### ERN1 is localized to the plasma membrane and the endoplasmic reticulum (ER)

The ERN sequence predicts several protein domains, including a transmembrane domain suggesting that ERN1 spans the plasma membrane. To determine the subcellular localization of ERN1, we transiently expressed ERN1 fused with a C-terminal m-Scarlet red fluorescent protein (ERN1-mScarlet) in *Nicotiana benthamiana* leaves. To label the plasma membrane, we transiently co-expressed LT16b, a plasma membrane localized marker, fused with N-terminal eGFP (eGFP-LT16b) (Cutler et al., 2000). Using confocal microscopy, ERN1-mScarlet was detected at the plasma membrane overlapping with eGFP-LT16b (Figure 4). To determine whether ERN1 localized to membranes other than the plasma membrane, we also transiently co-expressed ERN1-mScarlet with the ER retention four peptide sequence KDEL, N-terminally tagged with GFP (GFP-KDEL). ERN1-mScarlet was co-localized with GFP-KDEL indicating that ERN1 is also localized to the ER (Figure 4).

**Figure 4.**
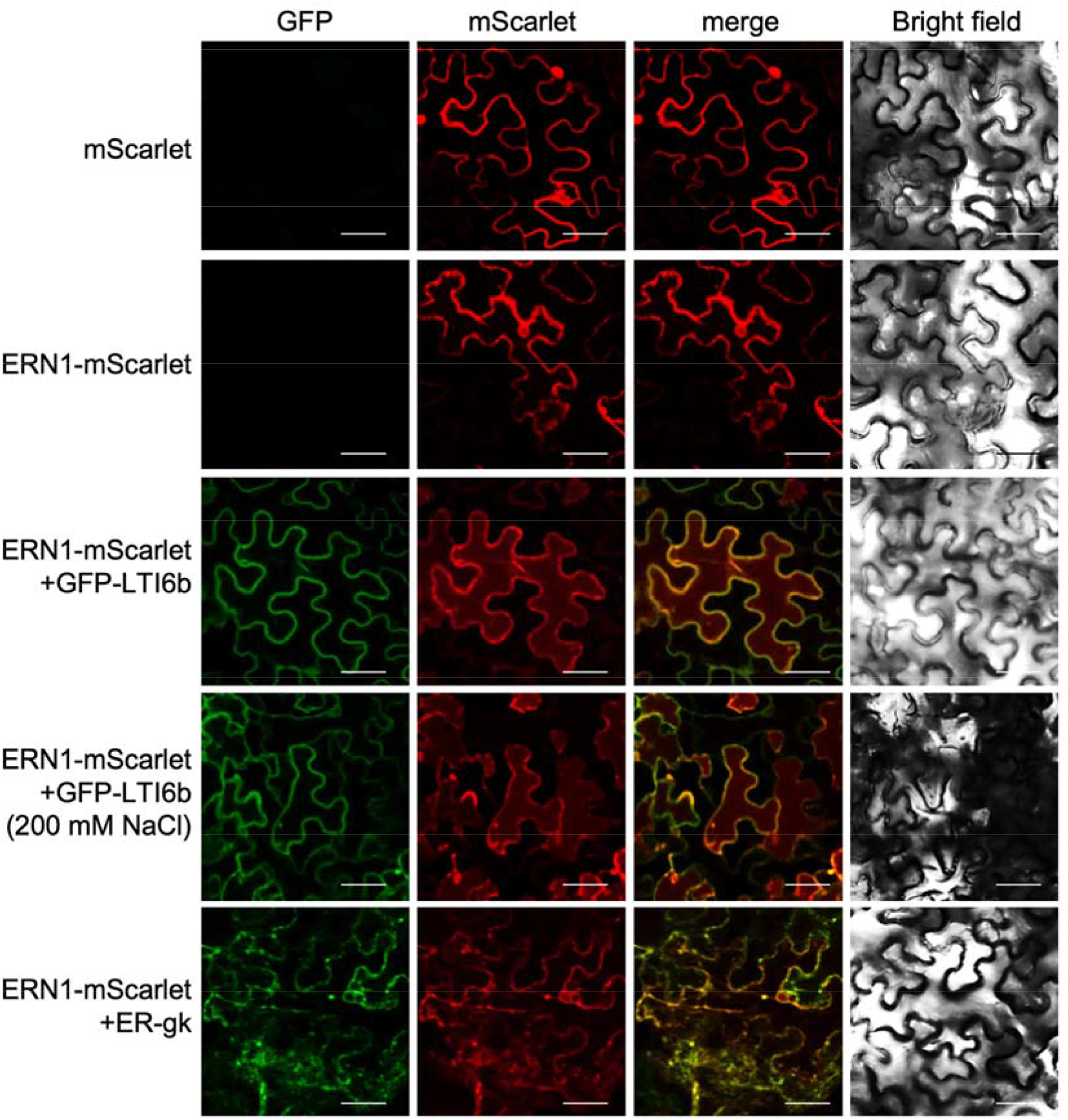
ERN1 is localized to the plasma membrane and ER in *Nicotiana benthamiana* leaf epidermal cells. mScarlet-tagged ERN1 (ERN1-mScarlet) was transiently co-expressed in *N. benthamiana* leaves with plasma membrane marker GFP-LTI6b and ER marker ER-gk, respectively. mScarlet and GFP signals were detected in leaf epidermal cells by a confocal microscope 48 h after infiltration with *Agrobacterium tumefaciens* expressing the different constructs. Scale bars=50 μm.

To confirm ERN1 subcellular localization, we generated transgenic *Arabidopsis*, stably expressing ERN1-GFP driven by the *ERN1* native promoter (*ERN1_pro_::ERN1-GFP*) in the *ern1.1* mutant background. Confocal microscopy of the transgenic roots confirmed the localization of the ERN1 to both the plasma membrane and the ER *in planta* (Figure 5). These results are consistent with ERN1’s function as a transmembrane receptor kinase.

**Figure 5.**
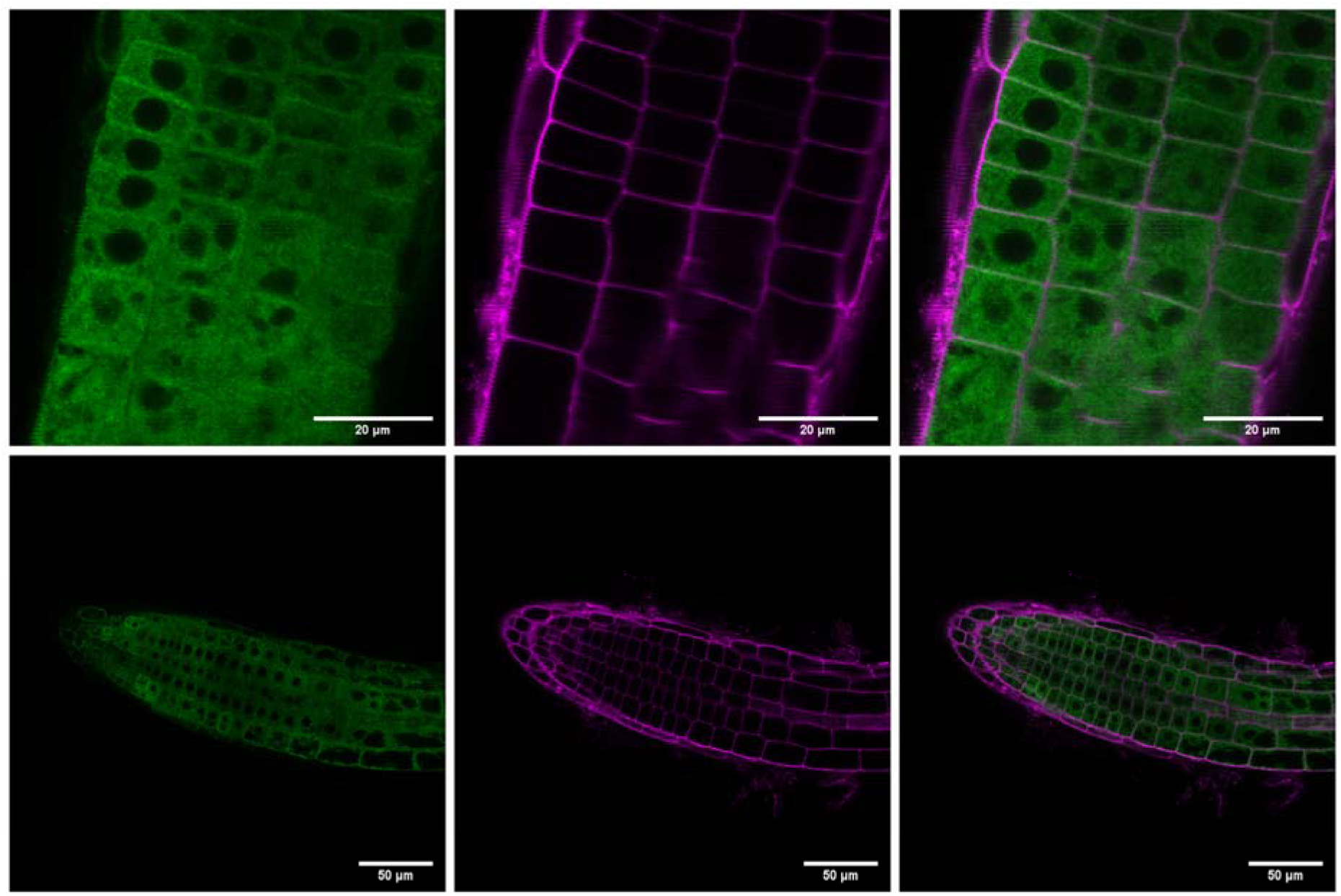
ERN1 is localized at the plasma membrane and ER in transgenic Arabidopsis roots. ERNI-eGFP fusion protein driven by ERN1 native promoter (ERN1_pro_ ::ERNI-eGFP) was stably expressed in *ern1.1* plants. Roots were stained with propidium iodide (PI) to visualize cell outlines and imaged with GFP fluorescence by confocal microscope. GFP alone (left panels), PI alone (center), and GFP+PI merged (right panels).

### Complementation of the *ern1.1* mutant results in enhanced plant growth and enhanced susceptibility to nematodes

To confirm that disrupt of *ERN1* by the T-DNA insertion is the cause of the enhanced nematode resistance phenotype, we complemented the *ern1.1* mutant with two different *ERN1* constructs, one driven by CaMV 35S promoter (*35S::ERN1*) and the other by its native promoter (*ERN1pro::ERN1*). Transgenic plants expressing of *ERN1* from its native or 35S promoter displayed enhanced plant growth phenotypes (Supplemental Figure S6 and Figure 6A). The enhanced growth phenotype was more profound at later stages of plant development (Figure 6A). To assess the nematode resistance phenotype, *ern1.1* plants complemented with both *35S::ERN1* and *ERN1pro::ERN1* were used in nematode infection assays. Our data show that lines expressing either transgene were significantly more susceptible to RKN than untransformed *ern1* mutants (Figure 6, B and C). In particular, the *35S::ERN1* complemented lines consistently displayed enhanced RKN susceptibility compared to WT, in further support of ERN1 function as a negative regulator of immunity (Figure 6B).

**Figure 6.**
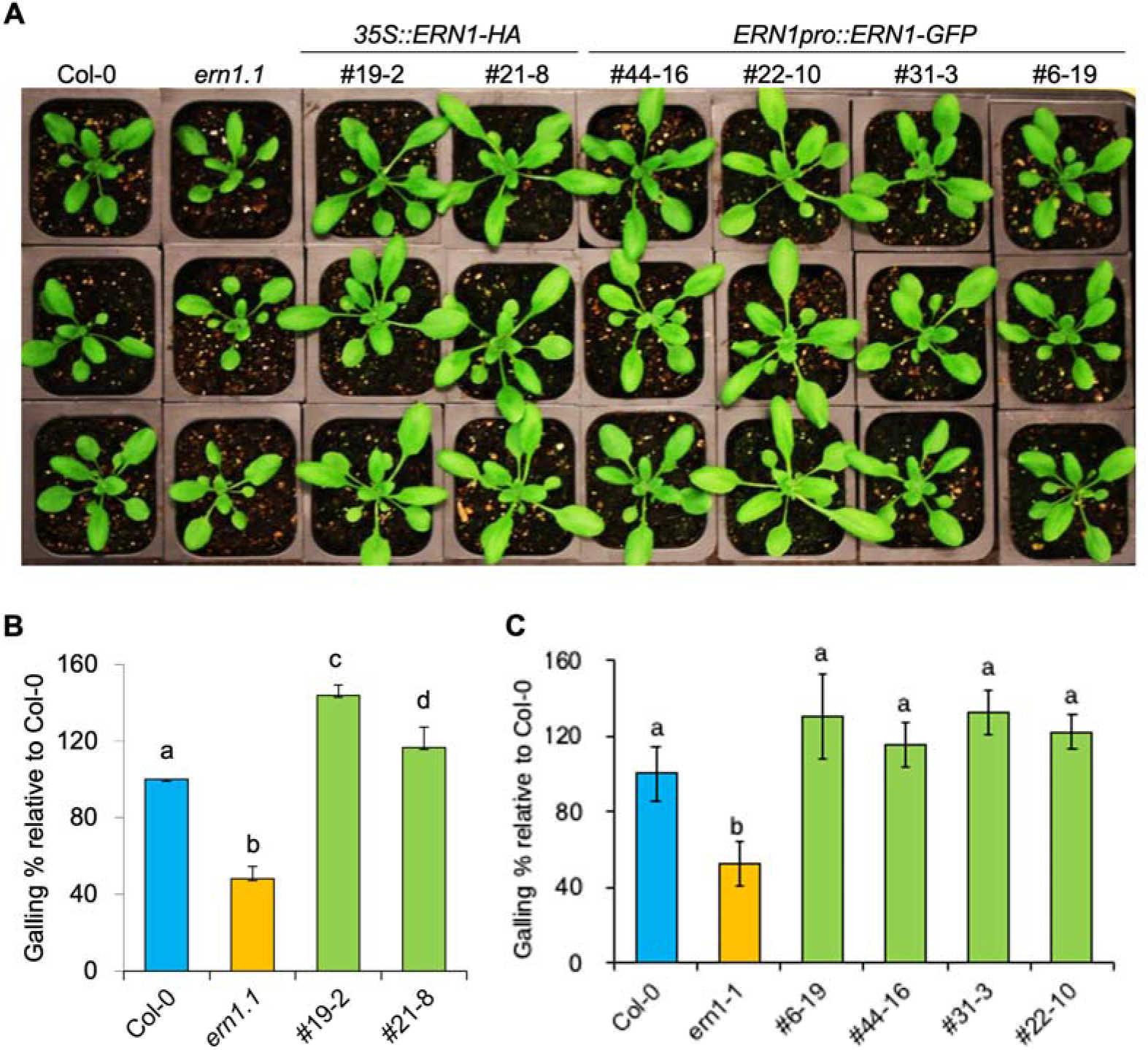
The *ern1.1* complemented plants exhibit enhanced growth and increased susceptibility to root-knot nematodes (RKN). (A) *35S::ERNl-HA* and *ERNlpro::ERNl-GFP ern1.1* complemented 5-week-old Arabidopsis lines grown at 12h photoperiod at 22’C. (B) RKN infection of *ern1.1* plants expressing *35S::ERNl-HA* and (C) *ERNlpro::ERNl-GFP* relative to Col-0. Arabidopsis plants grown on Gamborg agar plates were infected with 100 J2s and evaluated for galling 3-4 weeks later. Bars are means ± SE (n = 48 per genotype). Bars with different letters indicate significant differences (P <0.05, ANOVA).

### ERN1 is a negative regulator of immunity

The enhanced resistance displayed by the *ern1* mutants to RKN combined with enhanced defense marker analyses lead us to propose ERN1 acts as a negative regulator of immunity. To assess the role of *ERN1* in immunity, we evaluated the response of the *ern1* mutants to the well-known flg22 peptide elicitor by triggering a burst of reactive oxygen species (ROS). We found significantly higher levels of ROS burst in response to flg22 in *ern1.1* and *ern1.2* leaves compared to WT (Figure 7A and B). Because our nematode resistance phenotype was based on examination of roots, we also assessed H_2_O_2_ accumulation in roots upon flg22 treatment using DAB staining. Consistent with enhanced ROS burst in *ern1* leaves, darker staining was observed in the *ern1* mutant roots compared to WT (Figure 7C). Further confirmation of the role of ERN1 as a negative regulator of immunity was obtained by examining expression of the defense marker *MYB51* as triggered by flg22 treatment in leaves. Both *ern1* alleles displayed significantly higher *MYB51* transcript levels compared WT (Figure 7D). These results extend the function of ERN1 from resistance to nematode infection to a broader role in general pathogen response.

**Figure 7.**
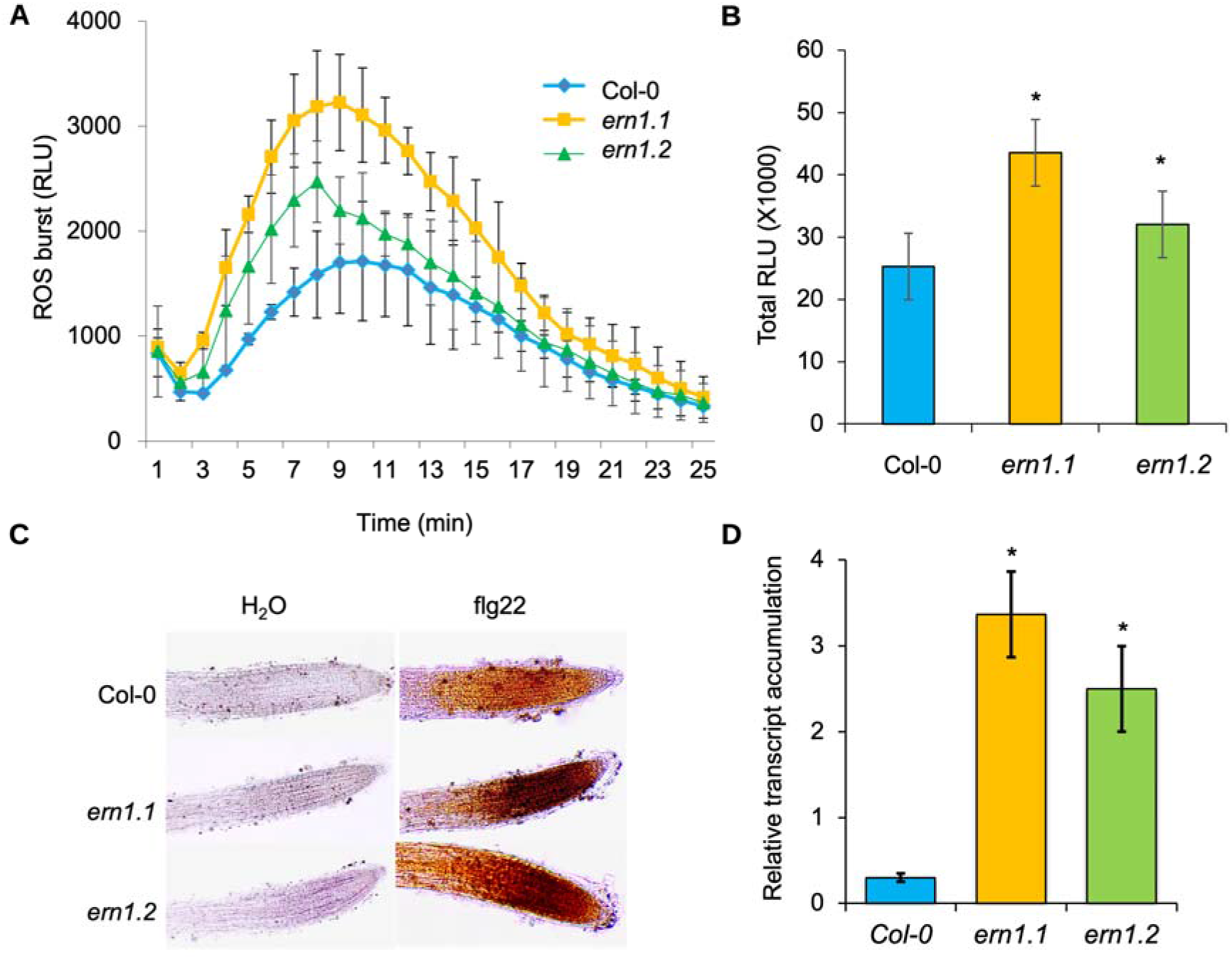
Flg22 triggered enhanced immune responses in the *ern* mutants. Flg22-induced reactive oxygen species (ROS) burst in leaves over time (A) or cumulative (B) and H_2_O_2_ accumulation in roots (C) of *ern1.1* and *ern1.2* mutants. (A and B) Leaf discs from 3-week-old plants were treated with 1 μM flg22 and the production of ROS was measured by a chemiluminescence assay and expressed as relative light units (RLU). Values are means + SD (n=4 per genotype). DAB staining (C) and *MYB51* expression by qPCR (D) in roots of 5-day-old Arabidopsis seedlings treated with 1 μM flg22 or water for 30 min. (D) *MYB51* expression was normalized to *UBQ22* and bars are means of three biological replicates ± SE. Asterisk denotes a significant difference (*P*<0.05, ANOVA).

Overexpression of *ERN1* in *ern1.1* (*35S:ERN1* lines #19-2 and #21-8) displayed significantly enhanced susceptibility to RKN compared to the *ern1.1* mutant as well as to WT. To further assess this putative reduced immunity, we examined these complemented lines for flg22-triggered ROS burst and H_2_0_2_ accumulation in leaves and roots, respectively. Both overexpression lines showed lower levels of ROS burst in leaves and less intense brown staining in roots compared to WT and *ern1.1* mutants, which is consistent with overexpression of *ERN1* leading to reduced immune responses (Figure 8).

**Figure 8.**
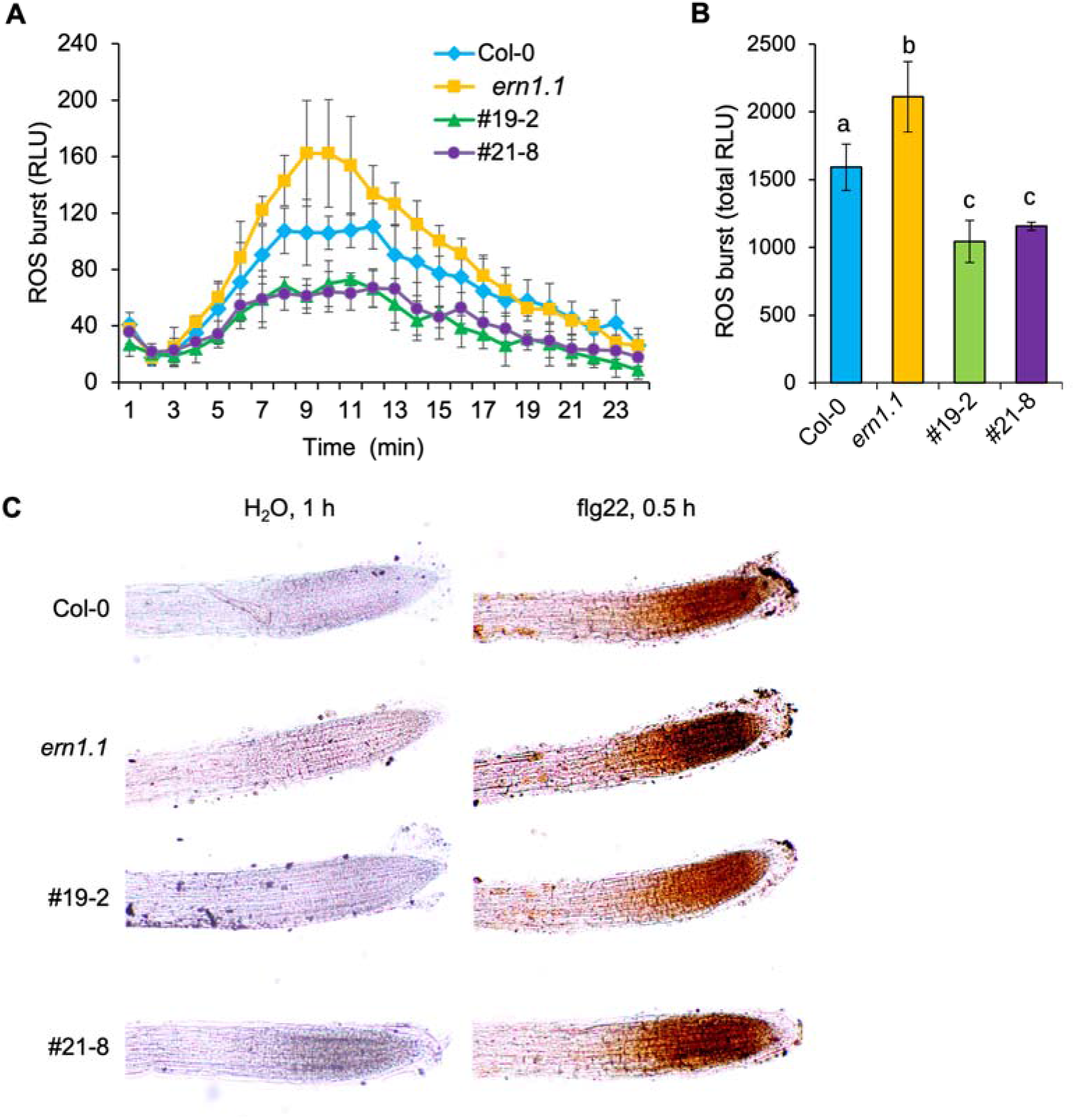
The *ern1* complemented lines have reduced levels of H_2_O_2_ accumulation in leaves and roots after flg22 elicitor treatment. (A) Time course and (B) total flg22-induced ROS burst in leaves of 3-week-old Arabidopsis *ern1.1* mutant and 35S::ERNl-HA *ern1.1* complemented lines (#19-2 and #21-8). Values are means ± SE (n=4 per genotype). Different letters indicate significant differences (P * 0.05, ANOVA). RLU= relative light units. C) DAB stained 5-day-old Arabidopsis roots treated with 1 μM flg22 for 30 min.

## DISCUSSION

Our quest to identify a cell surface localized nematode immune receptor unexpectedly resulted in identification of a negative regulator of plant immunity. We this immune regulator ERN1 and it is a G-LecRK belonging to a family with over 38 members in *Arabidopsis* (Teixeira et al., 2018). The G-LecRKs are proteins with an ectodomain that resembles the *Galanthus nivalis* agglutinin (GNA) mannose-binding motif and are also known as B-type LecRKs (Lannoo and Van Damme, 2014; Vaid et al., 2012). The best-known members of this group are the S-locus (S-locus glycoprotein/SLG containing) RKs, known for their role in self-incompatibility in flowering plants (Sherman-Broyles et al., 2007; Kusaba et al., 2001). While members of the L-LecRKs have been implicated in defense the G-type LecRKs have not been much explored for their role in immunity and defense (Sun et al., 2020). Our domain analysis identified six distinct domains (G-lectin, SLG, EGF, PAM, TM and kinase) in the ERN1 sequence, which is the largest number of domains identified among the members of this family. Besides lectin-binding and kinase functions, the cysteine-rich domain known as the EGF domain is thought to play a role in disulfide bond formation (Shiu and Bleecker, 2001), while the PAN motif specifies protein-protein or protein-carbohydrate interactions (Tordai et al., 1999). This suggests that ERN1 likely partners with multiple types of proteins to carry out its function.

Both *ern1* mutants displayed a reduced number of nematode infections and consequently enhanced resistance to RKN infection. Root growth and branching are important characteristics when it comes to RKN infection. This group of nematodes infect behind the root tips and the ability of roots to form branches with new root tips affects nematode infection. Our root growth measurements combined with lateral root capacity assays indicated that *ern1* mutants have similar root growth morphology as WT eliminating the possibility that the reduction in RKN infection rate is due to altered root formation. Normal root morphology, but an enhanced resistance phenotype suggests a specific role for ERN in immunity.

One of the challenges studying nematode immune responses, particularly the transiently induced responses, is synchronization of nematode penetration and infection without causing physical damage to the roots. To circumvent these challenges, nematode elicitors of immune responses have been identified, including NemaWater and infective-juvenile extracts (Teixeira et al., 2016; Mendy et al., 2017). Using RKN NemaWater, infective-juvenile extracts, and egg extracts we showed that several immune responses, including ROS bursts and defense gene expression are induced at higher levels in the *ern1* mutants. Preparation of NemaWater and infective-juvenile extracts require tedious and labor-intensive acquisition of clean infective-juveniles. In contrast, RKN eggs can be easily extracted from infected plants used to maintain the nematode cultures and are far more abundant than infective-stage juveniles. Therefore, our work also identified an simpler and more abundant source of RKN immune elicitors. It is likely that eggs and infective-stage juveniles have overlapping but also stage-specific immune elicitors. For example, nematode egg shells are highly enriched in chitin, while only trace amounts of chitin are present in the nematode juvenile esophageal lumen (Holbein et al., 2016).

It is well documented that immune responses are tightly regulated to protect against runaway immunity and its lethal consequence. Different mechanisms for immune regulation have been identified, including negative regulators of immunity (Lu et al., 2018). Our data indicates that ERN1 is a negative regulator of immunity. ERN1 localization at the plasma membrane and its effect on immune responses, induced by both nematode and microbial elicitors, indicate a wide role for this regulator in immunity against pathogens. How ERN1 suppresses immune responses remains unknown and requires further investigation. While *ern1* mutants exhibit enhanced immunity, we did not observe any necrotic or cell death-associated lesions, on above ground or below ground tissues, as seen on other enhanced immunity phenotypes also known as lesion mimic mutants (Dietrich et al., 1994; Greenburg et al., 1994; Weymann et al., 1995). Such lesions were also absent from the RKN infected plants indicating while *ern1* immune responses are enhanced, it is likely a balanced immune response. This observation also suggests that more powerful negative immune regulators are still working in the *ern1* mutant background. Furthermore, these results reveal that manipulation of an appropriate negative immune regulator may yield plants with desirable enhanced immunity without adverse effects.

The enhanced immune response in *ern1* mutants and absence of plant size and morphological phenotypes seem to contradict the expected balance between immune responses and plant growth. Elevated immune responses require more energy and are associated with a fitness cost. However, our observations do not detect a fitness cost in *ern1* mutants. Such fitness costs could exist but were not observed under our conditions or may be displayed in features we did not experimentally evaluate. Interestingly, while the *ern1* mutants were similar to WT, complementation of *ern1* mutants with an overexpression construct or with the native *ERN1* promoter yielded altered plant growth phenotype with both complemented lines having larger rosettes than the WT (Figure 6A). Despite this morphological similarity, complemented lines with the native promotor and WT had comparable levels of RKN susceptibility (Figure 6C), while the overexpression complemented lines exhibited higher levels of RKN susceptibility (Figure 6B). This enhanced susceptibility is associated with reduced levels of immune response (Figure 7) and further confirms the role of ERN1 as a negative regulator of immunity.

Several members of the G-LecRK family have been implicated in abiotic stress responses and evidence is starting to emerge as to their roles in biotic stresses (Sun et al., 2020). Both gene expression and mutant analyses indicate positive roles of G-LecRKs in immunity to microbial pathogens as well as to insect pests (Chen et al., 2006; Cheng et al., 2013; Gilardoni et al., 2011; Li et al., 2015; Ranf et al., 2015; Gouhier-Darimont et al., 2019; Sun et al., 2020). In contrast, here we report that the ERN1 G-LecRK is a negative regulator of immunity. Compared to *Arabidopsis*, the G-LecRK family has expanded in crops, such as tomato, and more than one orthologous sequence to *AtERN1* are identified in crops (Teixeira et al., 2018). It is important to determine whether the phenotype displayed in the *Arabidopsis ern1* mutant can be replicated in tomato. Editing these orthologous genes with CRISPR gene editing technology should be prioritized to potentially develop tomato plants with enhanced resistance to RKN. While resistance to RKN in tomato exists and is conferred by the *Mi*-*1* resistance gene, and appearance of resistance breaking RKN isolates are commonplace worldwide due to reliance on this single source of RKN resistance. Extending the insights gained here to *ERN1* homologs in crops may provide alternative sources of resistance to this harmful group of nematodes.

## Material and methods

### Nematode culture and inoculum preparation

*Meloidogyne incognita* isolate P77R3 was maintained on tomato cultivar Daniela (Hazera). Plants were grown in UC mix3 and sand (1:9, v/v), fertilized with MiracleGro® (Scotts Miracle-Gro Co) and kept in a glasshouse at 24° to 30°C.

Nematode eggs were extracted from roots using 10% bleach and sieving (Hussey and Barker, 1973). Eggs and plant debris collected on a 500 mesh sieve were fractionated twice on 35% sucrose and rinsed several times with sterile water. The eggs were sterilized by vortexing in 5% bleach for 5 minutes, rinsed several times with sterile water, re-sterilized by shaking in 50 mg/L nystatin and 30 mg/L gentamicin for 5 min and rinsed several times with sterile water. Sterilized eggs were either flash frozen in liquid N_2_ or hatched under sterile conditions using a modified Baermann funnel (Martinez de Ilarduya et al., 2001). Two days later, infective-stage juveniles (J2s) were collected, counted, and suspended in a 0.5% carboximethylcellulose solution.

### Plant material, growth conditions and nematode inoculation

*Arabidopsis thaliana* (Arabidopsis) Col-0 and mutants were obtained from the Arabidopsis Biological Resource Center (ABRC). Seeds were plated on ½ strength Murashige and Skoog (MS) basal salt media (Phytotechnology Laboratories) supplemented with 1% sucrose and 0.8% agar (Sigma-Aldrich) (pH 5.7) and maintained vertically in plant growth rooms with a 12 h photoperiod at 22°C. For RKN infection assays, seeds were plated on Gamborg media (Sigma-Aldrich) supplemented with 3% sucrose and 0.6% daishin agar (Bioworld) (pH 6.0) and maintained at 45° angle as described above. Two-week-old seedlings, with six seedlings per square plate, and 8-day-old seedlings, with 20 seedlings per square plate, were used for galling assay and RKN-induced gene expression analysis, respectively. Seedlings were inoculated with 100 J2s per seedling and plates were kept horizontally for 24 h in the dark. Later, plates were maintained as described above. For pathogenicity assays, galls on roots were evaluated four weeks after inoculation. To estimate the number of galls per root area, roots were photographed and surface area determined using ImageJ (https://imagej.nih.gov/ij/). The number of galls on Col-0 roots was defined as 100 percent and number of galls on mutant roots was reported relative to Col-0. RKN infection of Arabidopsis mutant lines were repeated once while infection of *ern1.1* (SALK_128729) and *ern1.2* (SAIL_63_G02) lines were repeated five times.

### Plant growth for phenotyping

Seeds for the different Arabidopsis genotypes were bulked at the same time for these experiments. For rosette leaf measurements, seeds were germinated on agar plates and 3-day-old seedlings were transplanted into UC soil mix 3 supplemented with Osmocote^R^. Plants were maintained at 12 h light photoperiod at 22°C, unless otherwise stated, for three weeks. Rosettes were photographed and measured using ImageJ.

Col-0 and *ern1* seedlings were grown side-by-side on agar plates for root length and lateral root capacity assays. For root growth rate, the position of the root tip was marked starting 4 days after placement in the growth room and marked 24 h later until 8 days. The plates were scanned and then lengths measured with ImageJ from the hypocotyl rootward for day 0-4 and between the consecutive marks for each time point thereafter. Lateral root capacity assays were performed as described in Moreno-Risueno et al. (2010). Briefly, plants were grown for 8 days then the root tips were excised, and plants were returned to grow for 3 more days. At 11 day, all emerged lateral roots were counted per root per genotype. This process prevents the formation of new lateral roots from the primary root and promotes the outgrowth of existing primordia allowing one to assess how many lateral roots had been specified at 8 days. Root growth rate and lateral root capacity experiments were performed twice.

### T-DNA insertion localization

All Arabidopsis mutant SALK, SAIL and GK lines were insertion mutants in Col-0 background. They were genotyped using both antibiotic selection and PCR amplification with gene-specific primers and T-DNA border primers (Supplemental Table S2). To determine the T-DNA insertion site in SALK_128729 and SAIL_63_G02 lines, homozygous plants were evaluated by PCR along with left or right border primers and a gene-specific primer in various combinations (Supplemental Table S2). Amplicons were sequenced and obtained sequences were aligned to At1g61550 sequence and to the T-DNA pROK2 and pCSA110 to precisely determine the T-DNA insertion sites in SALK_128729 (*ern1.1*) and SAIL_63_G02 (*ern1.2*) lines, respectively.

### Nematode juvenile/egg extracts and NemaWater

Frozen RKN J2s or eggs were ground in a mortar and pestle and the powder was suspended in PBS (pH 7.0) used immediately or frozen at −20°C overnight. The suspension was centrifuged at 9500 g for 15 min at 4°C and the supernatant was used to treat Arabidopsis seedlings or roots.

For NemaWater preparation, J2s were hatched in small volume of water for 48 h as described earlier (Mendy et al., 2017). The hatched J2s were spun down by centrifugation at 2000 rpm for 5 min and the supernatant was used as NemaWater.

### Gene expression analysis

RNA was isolated from Arabidopsis roots and leaves using GeneJET Plant RNA purification kit (Life Technologies) or Trizol (Life Technologies), respectively. Three μg of RNA was DNase-treated and used for cDNA synthesis using Superscript III reverse transcriptase enzyme (Invitrogen) and oligo-dT primers according to the manufacturer’s recommendations.

Quantitative real-time PCR analysis was performed using gene-specific primers (Supplemental Table S2) and iQ SYBR Green Supermix (Biorad) in iCycler5 IQ (Biorad) in 15 □l using the following program: 94°C for 5 min, followed by 40 cycles of 94°C for 30 sec, 58°C for 30 sec, 72°C for 30 sec and a final cycle of 72°C for 3 min. Three biological replicates, with two technical replicates each, were performed and the generated threshold cycle (ΔΔC_T_) was used to calculate transcript abundance relative to *UBQ22 or At*18S (Supplemental Table S2).

### Elicitor experiments and DAB staining

Five-day-old seedlings grown on mesh supported ½ MS plates, were placed into 6-well plates with liquid ½ MS media for 24 h. Then, they were exposed to egg extract or water, for 1 h or to 1μM flg22 for 30 min. The solutions were replaced with freshly prepared 1 mg mL^−1^ of DAB (3,3’-Diaminobenzidine; pH of 5.5) in 200 mM Na2HPO4 buffer for 45 min (Thordal-Christensen et al., 1997). Seedlings were washed three times with 50% (v/v) ethanol, mounted in 50% (v/v) glycerol and observed using a LEITZ DMRB Leica Microscope, objective 10X/0.65mm N Plan. These experiments were performed 2-3 times.

### Kinase activity

Roots of 12-day-old seedlings of wild type (WT; Col-0), *ern1.1* and *ern1.2* mutants, grown on ½ MS plates, were floated in water overnight. Then, the roots were treated with 5 eggs μL^−1^ egg extracts for 0.5 h, 1 h and 2 h or water control for 1 h. Protein extracts were fractionated onto 10% SDS-PAGE gel and activated MPKs were detected using anti-phospho-p42/44 MPK antibody. This experiment was performed twice.

### Vector constructs, complementation and microscopy

The *ERN1* overexpression complementation construct (35S:ERN1-3xHA), was generated in the Gateway vector pGWB14 containing a 35S promoter. The coding sequence of *ERN1* without the stop codon was amplified and cloned into pENTR221 and recombined into pGWB14 using a one-tube recombination protocol. Primers used for amplification and assembly of these constructs are listed in Supplemental Table S3. The *ERN1* native promoter driven complementation construct (ERN1pro::ERN1-mEGFP) was generated in a single reaction, using NEBuilder HiFi DNA Assembly Master Mix (NEB, E2621), to assemble three fragments into a modified pCAMBIA1300--MCS vector that lacks a promoter (Guo et al., 2021) (Supplemental Table S3). The promoter and coding sequence of *ERN1* without the stop codon was amplified from genomic DNA of WT Arabidopsis plants. Similarly, an overexpression *ERN1-mScarlet* construct was developed in *pCAMBIA1300-35S::ERN1-mScarlet*. All constructs were sequence verified before use.

Expression constructs were transformed into *ern1* mutant plants by the floral dip method (Clough and Bent, 1998) and transformants were identified using standard methods. For each reporter or complementation transgene, T2 lines that exhibited a 3:1 ratio of resistant:sensitive seedlings, indicating the transgene is inherited as a single locus were selected for propagation and homozygous T3 lines were used in further analyses. For each reporter, at least three independent lines were examined.

For subcellular localization in *N. benthamiana* leaves, *ERN1* was transiently expressed in 3-week-old plants by hand infiltrating *Agrobacterium tumefaciens* containing either one or two of the following constructs, *pCAMBIA1300-35S::mScarlet*, *pCAMBIA1300-35S::ERN1-mScarlet*, *35S::GFP-LTI6b* (Martiniere et al., 2012) or 35S::*ER-GFP* (ER-gk) (Nelson et al., 2007), using a needle-less syringe at a final OD600=0.5. Fluorescence in leaf tissues was detected after 48 h by a Leica SP5 confocal microscope. GFP and mScarlet, respectively, were excited at 488 and 543 nm, and images were collected at 498-530 and 555-635 nm. For the plasmolysis treatment, leaves were treated with 200 mM NaCl for 5~10 min before visualization.

For subcellular localization in Arabidopsis roots, stable transgenic T3 homozygous lines expressing *ERN1pro::ERN1-GFP* were grown on ½ MS plates. GFP fluorescence in 5-day-old roots was detected by a Leica SP8 confocal microscope using the settings described above.

### ROS burst assay

ROS burst in leaves was evaluated using 3-week-old Arabidopsis plants. Leaves were excised into 2 mm pieces using a blade and floated overnight on sterile water in a petri dish. Similar size leaves were transferred to a white 96-welll plate (Corning Costar) with 170 μl sterile water supplemented with 100 nM flg22, 20 μM luminol (Sigma), and 5 μg mL^−1^ horseradish peroxidase (Sigma). Luminescence was measured with a Tecan Infinite F200 plate reader. These experiments were performed 2-4 times.

### Statistical analyses

Statistical analyses were performed using GraphPad Prism (GraphPad Software, La Jolla, CA, USA). Comparisons were performed using Student *t*- test or One-way ANOVA and P < 0.05 were considered statistically significant.

## Supplemental Data

The following supplemental data are available in the online version of the article.

Supplemental **Figure S1.** *ERN1* genomic structure with locations of T-DNA insertions and gene expression in the *ern1* mutants.

Supplemental **Figure S2.** Shoot growth is not compromised in *ern1* mutants.

Supplemental **Figure S3.** Above ground growth is not compromised in the *ern1* mutants.

Supplemental **Figure S4.** Root growth is not compromised in *ern1* mutants.

Supplemental **Figure S5.** Nematode egg extract induces defense gene expression in Arabidopsis roots.

Supplemental **Figure S6.** Expression of *ERN1* and shoot phenotype of complemented plants.

Supplemental **Table S1.** List Arabidopsis RLK mutants screened with root-knot nematodes.

Supplemental **Table S2**. List of primers used in PCR and RT-qPCR.

Supplemental **Table S3.** List of primers used in cloning.

## Acknowledgements

We thank Professor Hailing Jin, UC Riverside, for helpful discussions.

